# Characterization of consensus operator site for *Streptococcus pneumoniae* copper repressor, CopY

**DOI:** 10.1101/676700

**Authors:** Henrik O’Brien, Joseph W. Alvin, Sanjay V. Menghani, Koenraad Van Doorslaer, Michael D. L. Johnson

## Abstract

Copper is broadly toxic to bacteria. As such, bacteria have evolved specialized copper export systems (*cop* operons) often consisting of a DNA-binding/copper-responsive regulator (which can be a repressor or activator), a copper chaperone, and a copper exporter. For those bacteria using DNA-binding copper repressors, few studies have examined the regulation of this operon regarding the operator DNA sequence needed for repression. In *Streptococcus pneumoniae* (the pneumococcus), CopY is the copper repressor for the *cop* operon. Previously, these homologs have been characterized to bind a 10-base consensus sequence T/GACAnnTGTA. Here, we bioinformatically and empirically characterize these operator sites across species using *S*. *pneumoniae* CopY as a guide for binding. By examining the 21-base repeat operators for the pneumococcal *cop* operon and comparing binding of recombinant CopY to this, and the operator sites found in *Enterococcus hirae*, we show using biolayer interferometry that the T/GACAnnTGTA sequence is essential to binding, but it is not sufficient. We determine a more comprehensive *S*. *pneumoniae* CopY operator sequence to be RnYKACAAATGTARnY (where “R” is purine, “Y” is pyrimidine, and “K” is either G or T) binding with an affinity of 28 nM. We further propose that the *cop* operon operator consensus site of pneumococcal homologs be RnYKACAnnYGTARnY. This study illustrates the necessity to explore bacterial operator sites further to better understand bacterial gene regulation.

## INTRODUCTION

Metals are essential nutrients to all living organisms. They are used as co-factors and structural components in a vast number of cellular processes. Iron, and manganese are examples of first-row divalent transition metals used by living organisms. Properties such as ability to form stable complexes play a vital role as to how each metal is used in the organism. The stability of biological complexes is characterized by the Irving-Williams series (Mn < Fe < Co < Ni < Cu > Zn) (1). In general, more stable complexes correlate to a metal’s toxicity, as native metals for an active site can be displaced by another metal ion further along in the observed series. This process is known as mismetallation and is primarily due to the promiscuity of different metal binding motifs (2). Higher order organisms have evolved ways to tightly regulate and use these metals, thus reducing some promiscuity in displacement. For most prokaryotes, however, metals like copper, nickel, and cobalt are broadly toxic. Within mammalian systems, copper is the most utilized and biologically relevant of these three metals (3-7). As such, in a process called nutritional immunity, mammalian hosts have evolved strategies to both sequester the universally necessary metals from bacteria (e.g., Fe, Mn, Ca) and bombard them with toxic metals such as copper and zinc (8, 9). Although Fenton chemistry mediated toxicity can occur in bacteria, the majority of copper-specific toxicity has been observed via mismetallation with iron-sulfur clusters, nucleotide synthesis, and glutamate synthesis (10-14).

Bacteria have evolved specialized import and export systems to acquire necessary metals and to adapt to metal toxicity. The presence of these import and export systems within the bacteria is usually based on need for the metal. Iron, for instance, is an essential metal for *Streptococcus pneumoniae* (the pneumococcus), a Gram-positive pathogen that causes pneumonia, meningitis, otitis media, and septicemia. In *S*. *pneumoniae*, iron has four known import systems, Pia, Piu, Pit, and the hemin binding system encoded by *SPD_1590* (D39 strain), but no known export systems (15, 16). Whereas calcium, zinc, and manganese all have export and import systems, the pneumococcus has no known import system for copper but contains a dedicated copper export system encoded by the *cop* operon (16-21). In general, Gram-positive, and some Gram-negative bacterial copper export systems consist of a *cop* operon regulator, a copper chaperone, and one or two copper exporters (17, 22-28). Mutations in the copper export protein in *cop* operons result in decreased bacterial virulence, highlighting the importance of nutritional immunity and copper toxicity in *S*. *pneumoniae* (17, 27, 29, 30).

The *cop* operon regulators function as either activators or repressors. Although there are *cop* operon activators and repressors in structurally distinct groups, they all serve to protect the bacteria against copper stress by sensing copper and facilitating its export. Activators are proteins that sense copper and activate gene expression in response, such as CueR in *Escherichia coli* (31). Occurring in species such as *Lactococcus lactis* and *S*. *pneumoniae* (CopR/Y), and in *Listeria monocytogenes* and *Mycobacterium tuberculosis* (CsoR), the *cop* operon repressors remain bound to DNA in environments lacking copper stress to block transcription, and release DNA upon binding copper (16, 24, 27, 32-34).

The CopR/Y family of *cop* operon repressors have consensus copper binding protein motifs, Cys-X-Cys. Each CxC motif (or CxxC) can bind in a 1:1 ratio with copper. These motifs can also bind zinc with a stoichiometry of two CxC motifs needed to bind one zinc (33). Copper binding causes a conformational change in the copper repressor leading to the DNA-binding release of what was thought to be the full *cop* operon operator, T/GACAnnTGTA (where n represents any nucleotide), while zinc binding leads to tighter *cop* operon operator binding (32, 33). However, the atomic and protein structural detail of how binding metal directly leads to the conformational changes associated with DNA binding is currently unknown.

Multiple studies regarding the *cop* operon have been performed in *S*. *pneumoniae* (11, 17, 27, 32, 33, 35, 36). The pneumococcal *cop* operon contains, *copY* as the repressor, *cupA* as a membrane-associated copper chaperone, and *copA* as the copper-specific exporter (16, 27, 32). Although the affinities of CopY/R family repressors for copper are generally high, the pneumococcal chaperone CupA is able to chelate copper from CopY, reduce it from Cu^2+^ to Cu^1+^, and transport copper to CopA for export (32, 35-37). CupA copper chelation allows for the recycling of CopY and its apo- or zinc-bound form to return to the *cop* operator to repress the operon. Pneumococcal CopY is homologous to several known antibiotic resistance repressors including BlaI, a *Staphylococcus aureus* MecI homolog that represses the gene for a β - lactamase (32, 33, 38). Like CopY, BlaI and MecI interact with a known operator sequence, TACA/TGTA, form a homodimer, and are mostly helical in secondary structure (32, 38). However, unlike CopY, BlaI does not have a known metal binding site and is regulated by proteases (39).

Here, we present bioinformatic data on the homology of *cop* operon operators and DNA-binding assays using recombinant pneumococcal CopY to characterize binding to the *cop* operon operator. We determined the consensus pneumococcal operator site, that pneumococcal CopY can bind to both operators relatively equally and independently, and that species with CopY homologs contain either one or two consensus operators but that species with two sites do not always have identical repeats.

## MATERIALS AND METHODS

### Aligning and Comparing CopY homologs and promoter sequences

The BLAST sequence alignment algorithm was used to align both *E*. *hirae* and TIGR4 *S*. *pneumoniae cop* operon promoter regions, the 21-base repeats upstream of the TIGR4 pneumococcal *cop* operon, and the promoter regions of pneumococcal species (40). A set of custom Python scripts (available from https://github.com/Van-Doorslaer/Alvin_et_al_2018) were used to assign identified *copY* homologs to bacterial genomes and extract the suspected regulatory region from individual species (100 bases upstream of the start codon). Importantly, in many cases, the initial blast search identified CopY homologs which matched multiple species’ isolates/strains. In this case, the identified proteins were again compared to the NCBI database, and the homolog with the lowest E-value was retained. The identified CopY homolog was not necessarily identical to the original query. If this approach was unsuccessful, the homolog was excluded from further analysis.

### Protein purification

CopY protein was purified as in Neubert et. al.(32), with modifications. The pMCSG7 vector includes an N-terminal 6x-His tag linked to CopY via a Tobacco Etch Virus protease (TEV) cleavage site(41). Unless specified, all steps were performed on ice or at 4 °C. After initial purification using immobilized metal-affinity chromatography (IMAC) (HisTrap FF, GE Healthcare), the crude CopY sample was incubated at 23 °C with a 100:1 mass ratio of recombinant TEV. The cleaved CopY was purified with subtractive IMAC (our TEV protease contains a C-terminal His tag). The flow-through was further purified by size-exclusion chromatography (SEC) (Superdex 200, GE Healthcare) using a buffer of 20 mM Tris pH 8, 200 mM NaCl, 1 mM tris(2-carboxyethyl)phosphine (TCEP). Peaks containing pure CopY (as determined by SDS-PAGE) were pooled and concentration was determined by absorbance at 280 nm. Samples were used immediately or protected with 35-50% glycerol and aliquoted into thin-walled PCR tubes containing 30 μL. These aliquots were then flash-frozen using liquid N_2_.

### Electromobility Shift Assay (EMSA)

Primers for binding, 5’-TAATTGACAAATGTAGATTTTAAGAGTATACTGATGAGTGTAATTGACAAATGTAGATTTT -3’ and 5’ – AAAATCTACATTTGTCAATTACACTCATCAGTATACTCTTAAAATCTACATTTGTCAATTA – 3’, were annealed by heating a 1:1 molar equivalent of each strand to 95°C, then reducing the temperature by ∼1°C/minute to 22°C. EMSA buffer was Tris-borate (TB) electrophoresis buffer (EDTA was left out to diminish metal chelation). Samples were incubated at 4°C for 5 minutes, loaded onto a 5% polyacrylamideTBE gel (Bio-Rad) that had been pre-run for 15 minutes in TB Buffer. Samples were electrophoresed at 40 V for 120 minutes. The polyacrylamide gel was stained with 0.02% ethidium bromide (Amresco) and imaged a Gel Doc XR+ System (Bio-Rad).

### Octet DNA/protein binding

Double stranded DNA fragments were prepared by incubating a 5’ biotinylated ssDNA with a complementary strand at 95 °C for 5-10 minutes then left on benchtop to cool to room temperature (Table S1). The dissociation constant for CopY with various DNA fragments was determined using an Octet Red384 (Pall ForteBio). Streptavidin biosensors (Pall ForteBio) were hydrated at 26 °C using the Sidekick shaker accessory for 10 min at 1000 rpm. Biotinylated DNA fragments were diluted to 250 or 50 nM, depending on the levels of biosensor loading, in the assay buffer (50 mM Tris pH 7.4, 150 mM NaCl, 4% glycerol, 1 mM TCEP, 1 mM NaN_3_, 0.1% Bovine Serum Albumin (BSA). During initial optimization, we observed significant non-specific binding of CopY to the biosensors in the absence of BSA. The inclusion of 0.1% BSA eliminated signals of nonspecific CopY binding at the highest concentrations used in the assay (3 µM). Hydrated sensors were incubated in the assay buffer to acquire a primary baseline. The sensors were then loaded with biotinylated dsDNA, followed by a secondary baseline measurement using wells with buffer solution. DNA-loaded biosensors were then moved to wells containing varying CopY concentrations to measure association then placed back into assay buffer for dissociation recordings. All experiments were maintained at 26 °C with shaking at 1000 rpm. The optimized protocol was as follows: 1° baseline 60 s, DNA loading 300 s, 2° baseline 180 s, association 180 s, and dissociation 270 s. The methods were optimized to minimize sharp association peaks and minimize recording at equilibria.

Analysis was performed using the Octet software. We applied a 1:1 (for dsDNA including one site) binding model using a global fit to biosensor replicates at each concentration of CopY. During pre-processing, an average of the 2° baseline across the various biosensors was applied, as well as Savitzkty-Golay filtering to reduce noise. The data were inter-step corrected using an alignment to the dissociation step. Data were modeled using combined fits of k_a_ and k_d_ values across independent replicates. Final estimates for K_d_ and related statistics were taken from the kinetic analysis.

Rate constants for each sample were determined using the Octet analysis software as follows. For all 1:1 stoichiometric modelling, complex formation was evaluated as pseudo-first-order kinetics. The observed rate constant (k_obs_) was calculated according to the equation 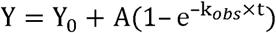. Where Y_0_= initial binding, Y = level of binding, t = time, and A = asymptote value at max response. Dissociation rate (k_*d*_) was calculated according to the equation 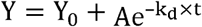. The calculated k_*obs*_ and k_*d*_ values were then used to determine k using the equation 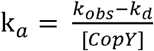 Finally, the dissociation constant (K_d_) was determined by the identity 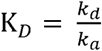

## RESULTS

### *cop* operator homology and frequency

Early DNA-binding studies were carried out using a CopY homolog from *Enterococcus hirae* on the interactions with the *cop* operon operator (22, 42, 43). We recently observed that there are two large repeats upstream of the pneumococcal *cop* operon that include a 10-base sequence important for CopY binding. These motifs differed slightly from those observed in *E*. *hirae* (Figure 1A). Although the amino acid sequence of the *E*. *hirae* copper repressor and the upstream binding repeats are highly similar to *S*. *pneumoniae* (32, 33), and contain the 10-base sequence, *E*. *hirae* operators upon initial observation lacked the extended regions flanking this sequence in pneumococcus (Figure 1B) (22). A BLAST search revealed that the 61-base stretch of DNA upstream of the pneumococcal *cop* operon that includes the two extended 21-base repeats is highly conserved in all pneumococcal species (40). Related searches identified that some bacterial species only contain a single operator upstream of their respective *cop* operons. DNA-binding studies for the *cop* operon operator and CopY/R (including homologs) have not been performed for two identical operators to determine *in vitro* affinity values with the closest being Portmann et. al (42).

**Figure 1.**
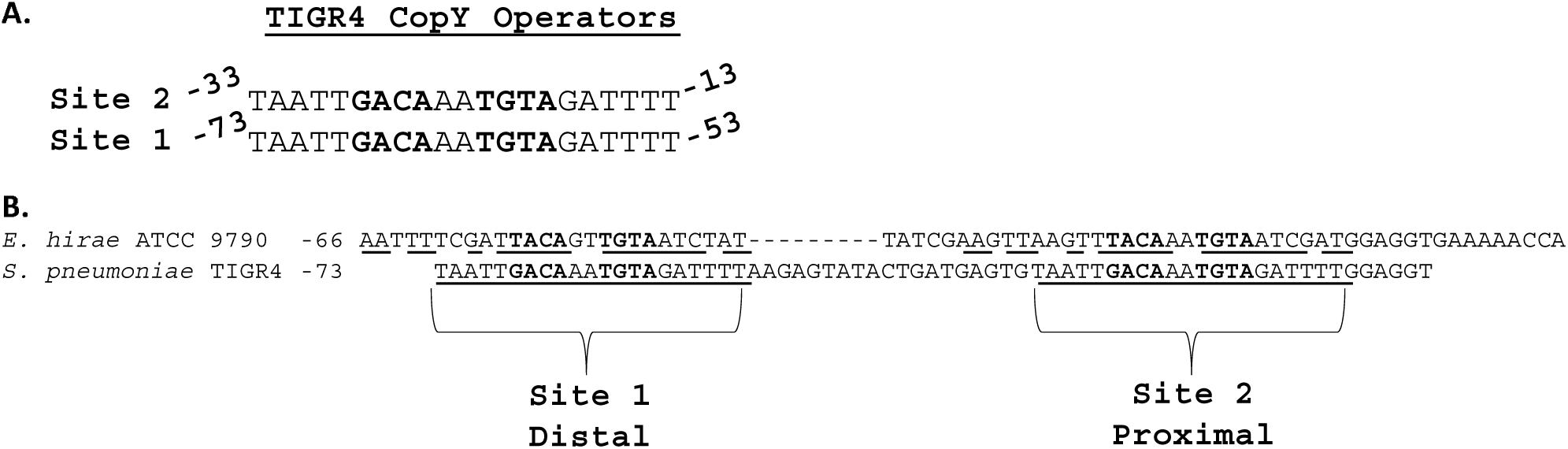
TIGR4 has two 21-base repeats containing the consensus CopY operators. **A.** Aligned 21-base sequences for the two CopY operators. **B.** TIGR4 *SP_0727* promoter region sequence (containing both 21-base repeats) aligned with the *E*. *hirae* ATCC strain 9790. Identical bases are underlined for the respective regions containing the operator.

We performed BLASTp searches for *S*. *pneumoniae* TIGR4 CopY homologs first excluding, then specific to the *Streptococcus* genus (40). Using a max target sequence number of 1000 for each search, then combining both lists, we found 335 different entries (Table S2).

From this list, we extracted protein sequences from the unique NCBI accession numbers (many NCBI accession numbers represented several/identical species) and then, the 100 bases upstream of the respective copper repressor start codon (Table S3, S4). Many of the 141 unique protein sequences belonged to species in the *Streptococcus* and *Lactobacillus* genera (Table S3). This table also included species such as the yogurt probiotic *Lactobacillus acidophilus* and *Mycobacteroides abscessus*, an emerging multidrug resistant pathogen that causes lung, skin, and soft tissue infections) (44).

We used programs within the MEME suite to identify DNA repeats within a 100-base upstream fragment, which would correspond to promoter and suspected operator-containing regions from 88 unique sequences (45, 46). As predicted by known CopY/R operators, the dominant 21-base sequence contained T/GACAnnTGTA (hereafter KACAnnTGTA) (Figure 2A) (47). Of these 88 sequences, 67 had two CopY/R *cop* operon operators such as TIGR4, *Streptococcus pyogenes, E*. *hirae*, and *Enterococcus faecium*, 14 had one *cop* operon operator such as *Enterococcus faecalis*, and 7 had none with no other consensus sequence found (Figures 2B-D, Tables S5-S7). Increasing the bases upstream maximum to 500 did not yield additional sequences (data not shown). For the genomes with two *cop* operators, we found that most of the operators were between 24 and 39 bases from each other with the mode being 26 bases (Figure S1). Of the species that had a single *cop* operon operator, the consensus sequences were more variable KACAnn<u>Y</u>GTA (Table S6). Of these 14 sequences, 7 were on the positive strand (but six were palindromic) and 7 non-palindromic sequences were on the negative strand (Table S6).

**Figure 2.**
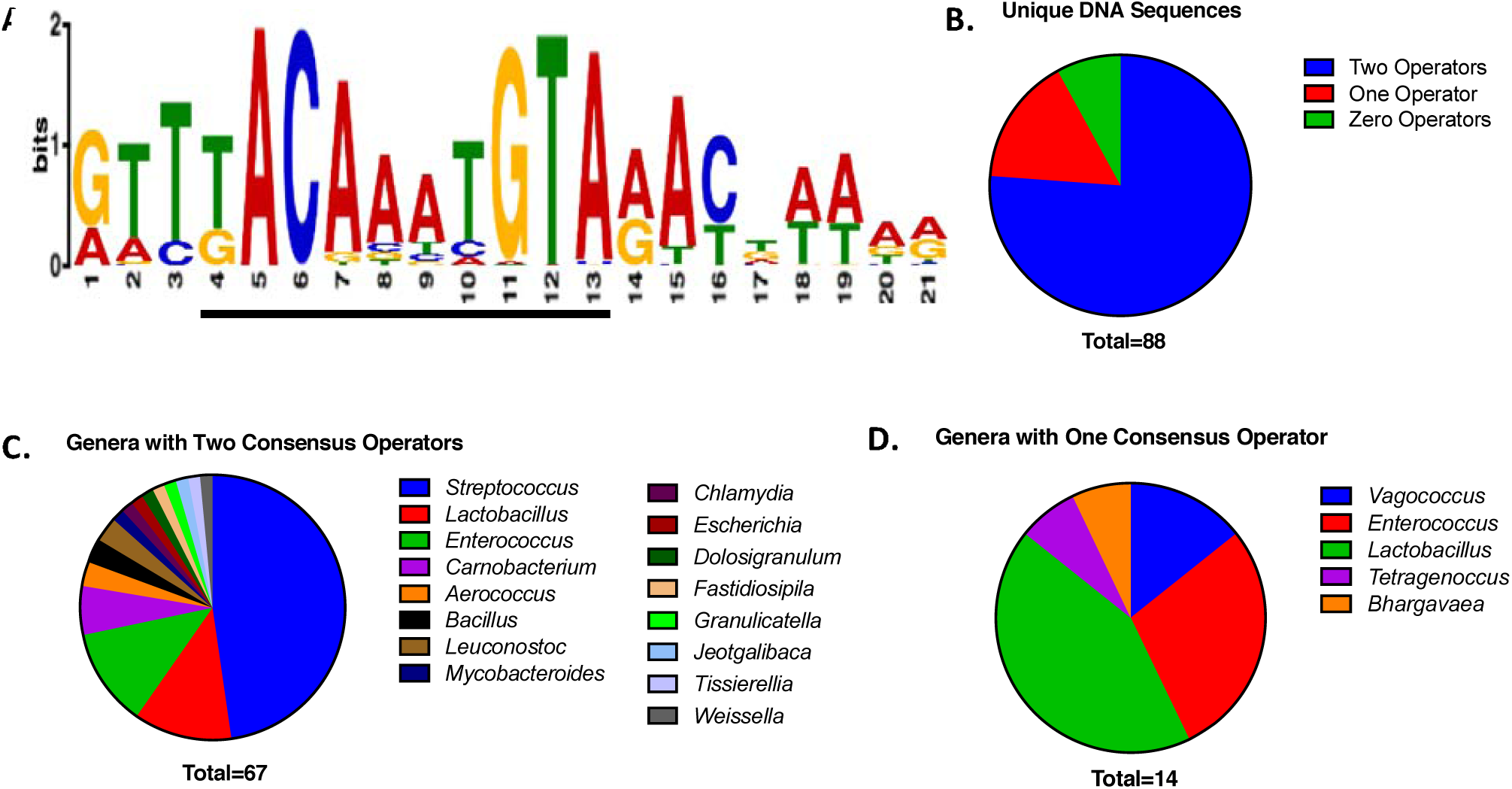
CopY *cop* operon operator consensus sequences in the genomic DNA. **A.** Top consensus DNA sequence contained within the 87 unique upstream 100 bases sequences to the respective CopY organism as detected by MEME suite. The literature-based T/GACAnnTGTA is underlined within the program generated consensus sequence. **B.** Total unique DNA sequences from Table S5 listed by number of CopY operators. **C.** Unique genera with two CopY *cop* operon operators from Tables S5 and S6. **D.** Unique genera with one CopY operator from Tables S5.

With the sequences that had no consensus *cop* operon operators for CopY/R, six were in the *Lactobacillus* genus and one in the *Macrococcus* genus (Table S7). For these seven strains that had no consensus *cop* operon operator, MEME suite did not identify a consensus sequences consistent across those seven strains. A BLAST search also found no significant similarity between the DNA fragments. Additionally, a separate, *in silico* search of these operators did not yield a consensus sequence consistent between the strains.

### CopY binds to both *cop* operators

Previous studies showed that CopY specifically bound to the *cop* operon operator in a sequence and metal specific manner as disrupting the operator bases or adding copper disrupted CopY binding, while adding manganese or iron had no detectable effect (32). These studies were done with only one full operator intact (32, 33). Thus, using an electric mobility shift assay (EMSA), we qualitatively tested CopY binding to DNA to the two-operator 61-base dsDNA fragment. As evaluated by EMSA, CopY bound in a dose-dependent manner (Figure 3).

**Figure 3.**
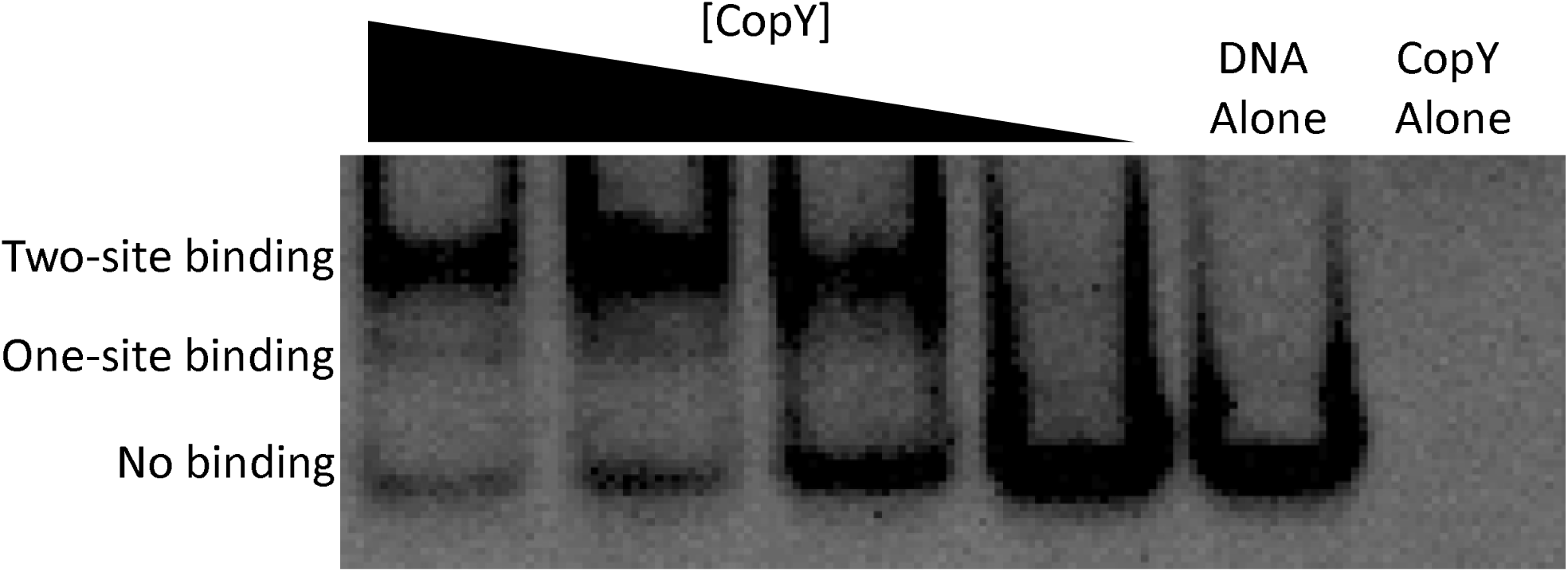
CopY binds to both *cop* operon operators. EMSA with CopY and two-site (operator) DNA. In seven of eight wells, a final concentration of 50 nM DNA was used with protein concentrations titrated by 2.5-fold dilutions from 640 nM to 41 nM. A final concentration of 640 nM CopY was used in a protein-without-DNA control with each replicate.

Consistent with having two different *cop* operon operators, titrating CopY with the two-operator DNA showed two distinct shifts via EMSA (Figure 3).

To quantitatively determine affinities to one or both operators, we used biotinylated DNA oligos and recombinant CopY protein using biolayer interferometry (BLI). All BLI binding experiments are done with dsDNA unless otherwise noted. These experiments demonstrated that the two-site DNA had a similar affinity (K_d_ = 28.1 nM) to the proximal DNA (DNA that has the distal site scrambled) (K_d_ = 25.5 nM) (Table 1, Figure 4 A, B). The distal site DNA (DNA that has the proximal site scrambled) had a slightly lower affinity (K_d_ = 55.2 nM) (Table 1, Figure 4B). However, a 2-fold change in affinity is within the error of the machine, thus indicating that all sites bind similarly and independently of each other. Further, this 2-fold difference could be attributed to the biotin tag on the 5’ end of the DNA slightly interfering with binding. Given the comparable affinities, it is likely that binding at each operator is non-cooperative—i.e., CopY dimers do not appear to have quaternary. As expected, CopY bound DNA constructs containing intact 21-base repeat containing *cop* operon operators with significantly higher affinities compared to a scrambled DNA negative control (scram) or a ssDNA containing the two-site operator which both exhibited extremely weak binding (Figure 4D, data not shown). Taken together, CopY binds both 21-base repeats containing the operator sites independently of each other with nanomolar affinity.

**Table 1.** Data and model statistics from Octet kinetic experiments. Listed dsDNA fragments and conditions were used with streptavidin probes and tested with recombinant CopY protein.

| Construct | K<sub>d</sub> (nM) | R<sup>2</sup> |
| --- | --- | --- |
| Two-Site | 28.1 ± 0.2 | 0.9474 |
| Distal Site | 55.2 ± 0.4 | 0.9432 |
| Proximal Site | 25.5 ± 0.2 | 0.9475 |
| 16-base | 360 ± 4 | 0.9808 |
| 19-base | 37.1 ± 0.3 | 0.9552 |
| 19-base T to C | 40.4 ± 0.3 | 0.9408 |
| *E. hirae* (Proximal) | 164 ± 2 | 0.9269 |

**Figure 4.**
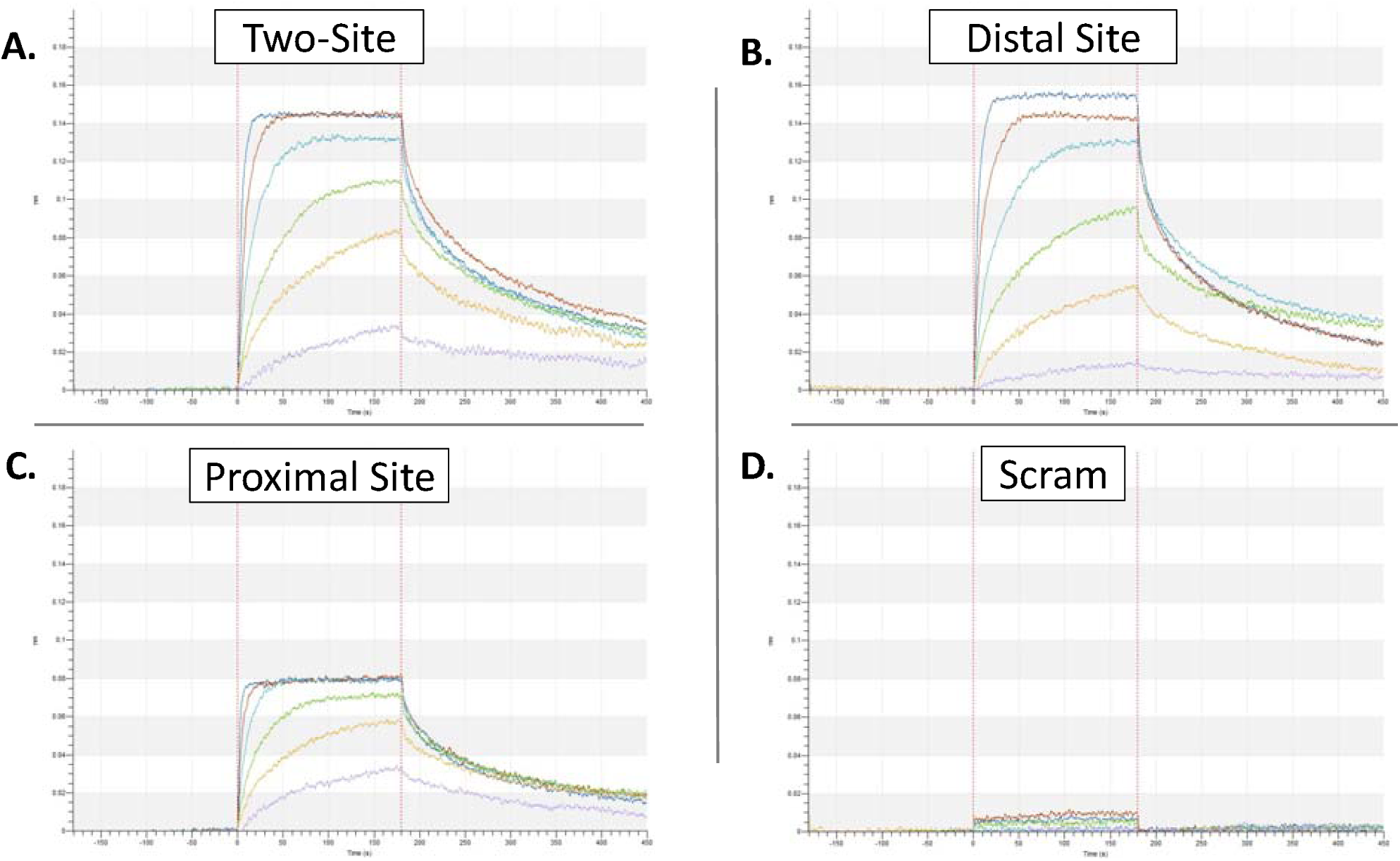
Affinity measurements for CopY and the *cop* operon operators. DNA fragments were loaded onto a biosensor and tested with 1000 nM (blue), 500 nM (red), 250 nM (light blue), 125 nM (green), 62.5 nM (orange), 15.6 nM (purple) CopY (A-D). **A.** Two-site **B.** Distal site **C.** Proximal site **D.** Scram. For each figure, data is representative of at least three experimental replicates.

### Determining a consensus CopY-family operator

With a reported consensus binding sequence of KACAnnTGTA on the leading strand, we hypothesized that CopY may also bind other similar sequences as previously seen in *Lactococcus lactis* (48). Allowing for one base variation from the reported binding sequence, we found matches upstream of genes upregulated under copper stress and hypothesized that they may also be regulated by CopY (11, 49). Seven potential binding sites were assessed using BLI. To our surprise, CopY did not bind to any of the fragments (Figure 5, Table 2, S2).

**Table 2.**
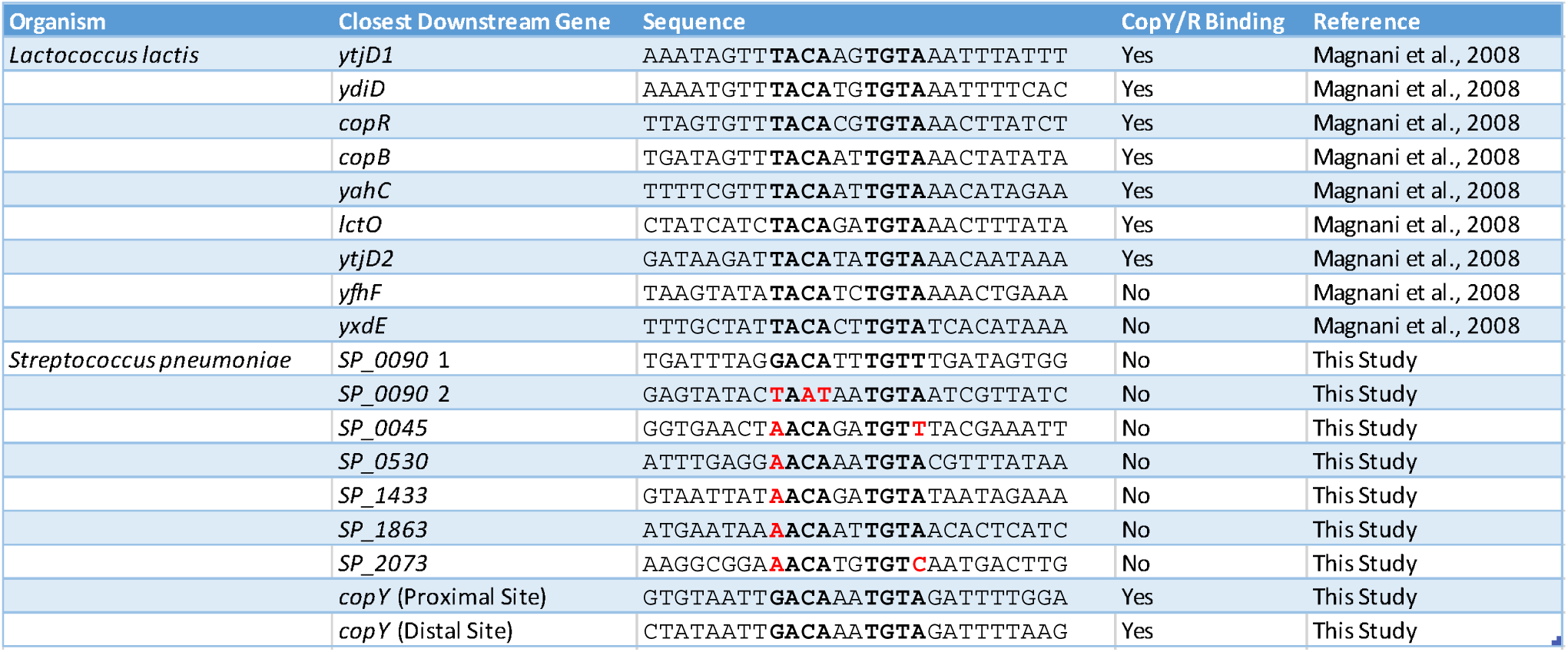
Outcomes of CopR or CopY binding to potential operator sites from *L*. *lactis* (Magnani et al., 2008) or *S*. *pneumoniae* respectively. Red bases indicate bases varying from the reported 10-base consensus sequence.

**Figure 5.**
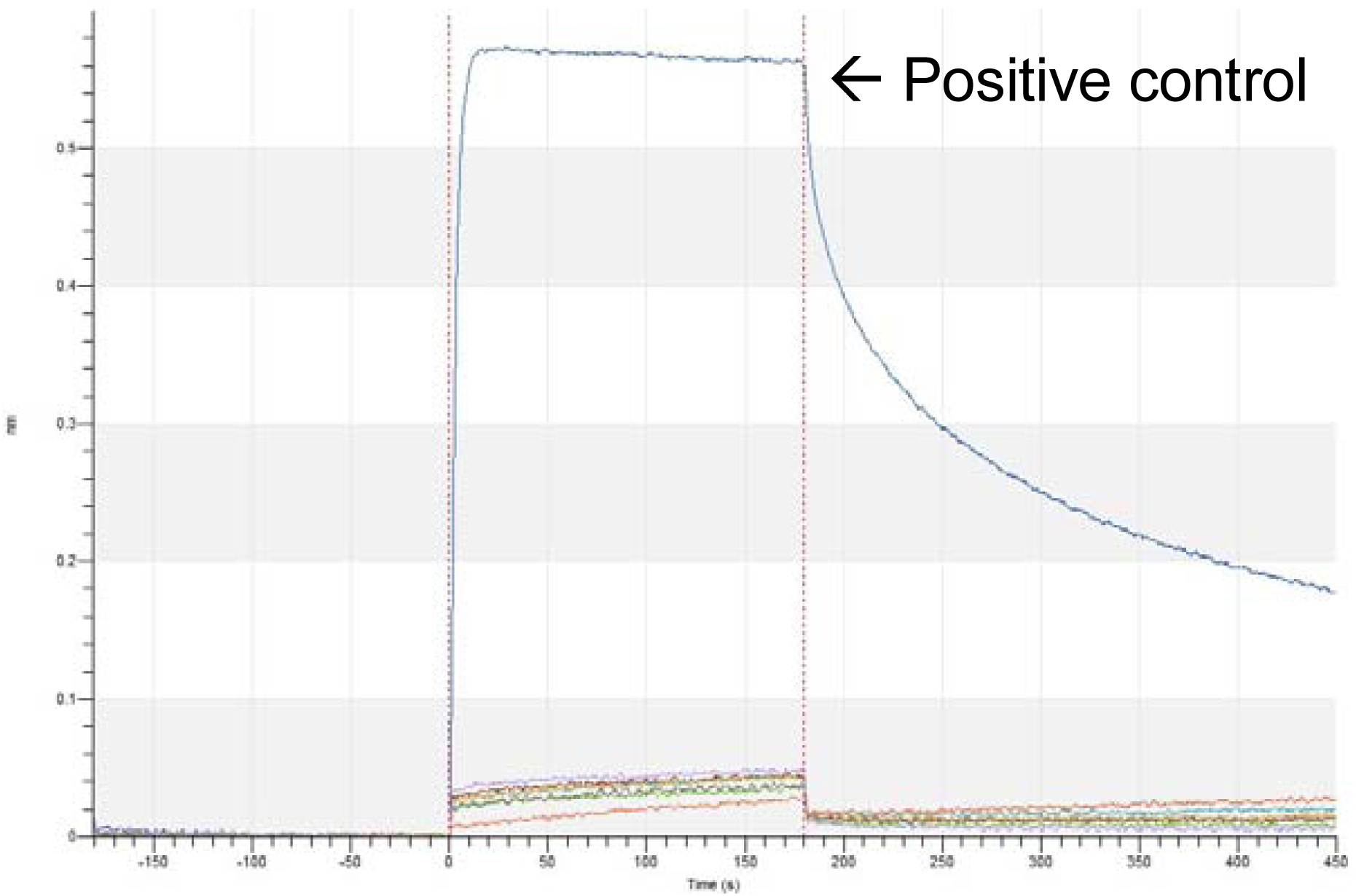
Prediction of CopY binding based on 10-base sequence overestimates binding sites. CopY at 3 µM was used to assess binding to DNA fragments containing potential CopY operators upstream of the respective genes *SP_0090* 1 (red), *SP_0090* 2 (light blue), *SP_0045* (green), *SP_0530* (orange), *SP_1433* (purple), with controls for two-site (blue), scram (gray), no DNA (red-orange). For each figure, data is representative of three experimental replicates.

Based on this result we suspected that the reported consensus sequence in the literature may be necessary, but not sufficient for CopY binding. For this experiment, we took a scrambled negative control DNA and added back only the known 10-base consensus sequence to where it exists in the second operator site. We found that CopY did not bind this fragment above the level of our negative controls (Figure 6A). Taken together, these and with prior data, we have concluded that the reported consensus binding operator is necessary, but not sufficient for binding (22).

**Figure 6.**
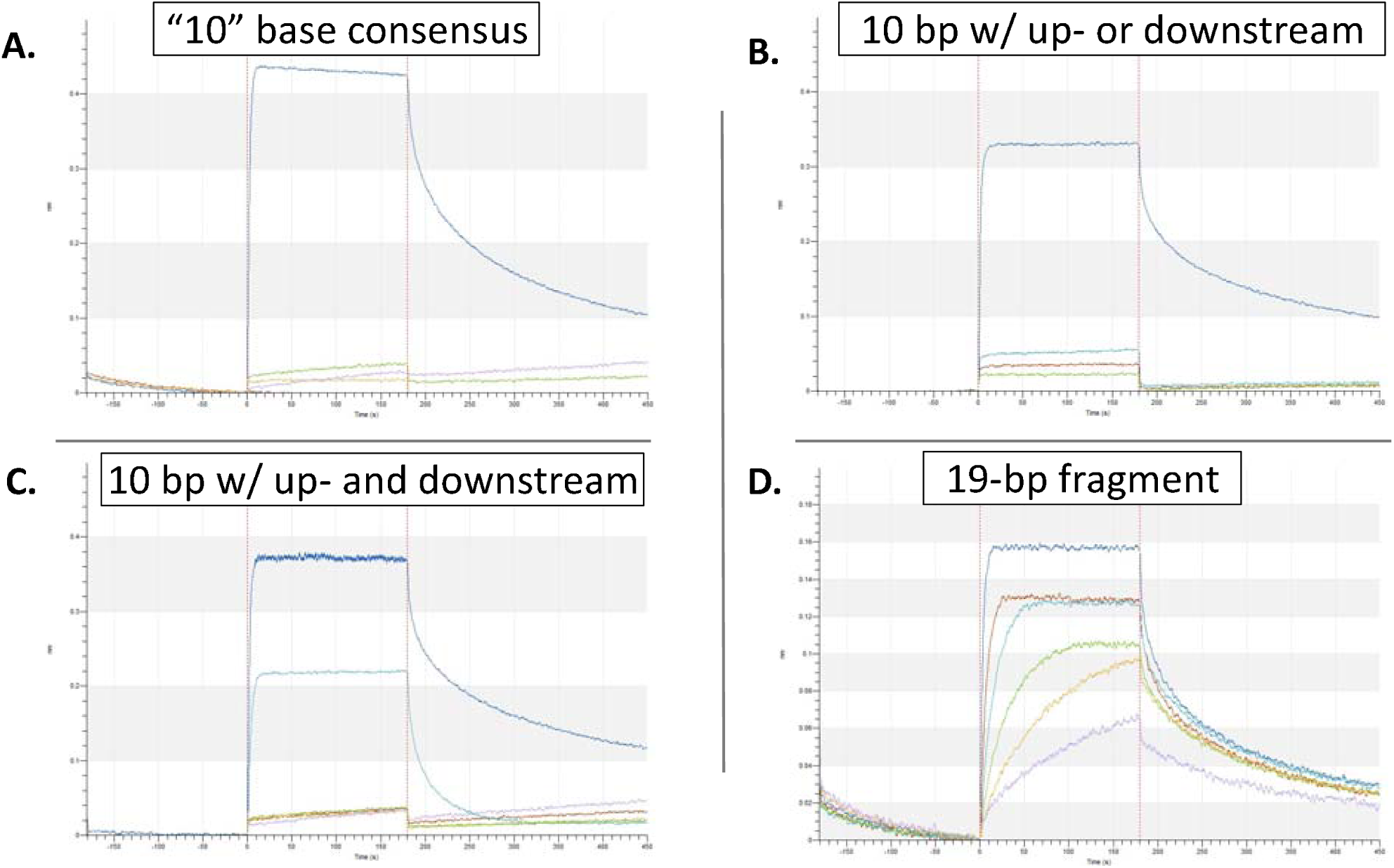
The minimum operator for sufficient CopY binding is 16 bases in length. **A.** Binding of the 10-base fragment compared to positive and negative controls. Two-site (blue), Two-site no protein (red), 10-base (green), scram (orange), no DNA (purple). **B.** DNA fragments containing either the upstream (5 bases) or downstream (6 bases) of the 10-base sequence within 21-base repeat compared to positive and negative controls. Proximal site (blue), five bases upstream (red), six bases downstream (light blue), 10-base sequence (green). **C.** Fragments were used to assess extended sequence on both sides of the 10-base sequence. Proximal site (blue), 14 bases (red), 16 bases (light blue), 10 bases (green), scram (orange), no DNA (purple). Panels A-C used 3µM CopY to assess binding to each of the fragments. **D.** A fragment containing 19 of the 21 bases in the repeat has comparable levels of binding to the fully repeat. The following concentrations were used to establish the K_d,_ 1000 nM (blue), 500 nM (red), 250 nM (light blue), 125 nM (green), 62.5 nM (orange), 15.6 nM (purple) CopY. Data is representative of three experimental replicates.

We next wanted to establish which bases outside of the previously reported 10-base consensus sequence were necessary for CopY binding. Using the proximal 21-base motif as a model, two fragments were generated: one containing the five bases upstream and another containing six bases downstream of the sequence. Neither of these fragments showed notable binding above that of the scrambled constructs suggesting that there are important bases on both sides of the 10-base sequence (Figure 6B). Next, two new constructs were tested to assess binding; one had the 10-base consensus sequence + two bases upstream and downstream for a total of 14 bases out of the full 21-base motif, and one had the 10-base consensus site + three bases to each side of the consensus was created for a total length of 16 bases out of the full 21-base motif. The 14-base sequence did not display binding above that of negative controls (K_d_ > 3 µM), but the 16-base sequence displayed binding (Figure 6C, S3, Table 1). While these 16-base DNA had a ∼10-fold lower binding affinity than the full 21-base site, these levels of binding suggest that the 16 bases make up the core operator site recognized by CopY and additional bases increase affinity at the site. Extending the sequence to 19 of the 21 bases led to comparable levels of binding to the full sequence (Figure 6D).

Taking into account 1) our results for adding three bases on each side of the 10-base reported sequence in pneumococcus, 2) looking externally of the 10-base fragments in *L*. *lactis* that it was predicted the repressor would bind to (with some binding and some not binding), and 3) analyzing the alignment of bacterial species operator sites via Meme Suite (Figs. 1A, 2A, Table S5) a clear pyrimidine-n-purine motif on each side of the 10-base sequence was revealed to be RnYKACAnnTGTARnY (where “R” is purine, “Y” is pyrimidine, and “K” is either G or T) (48).

The initial characterization of the *cop* operon operator site occurred using the *E*. *hirae* genome (22, 50). Thus, to gather more details on the consensus pneumococcal CopY operator site, and test this extended operator hypothesis, we used *E*. *hirae* DNA with the distal or proximal operators intact and examined it for pneumococcal CopY binding. The *E*. *hirae* DNA had several differences to test the new operator hypothesis; 1) the *E*. *hirae* DNA sites have a “T” instead of the G found in *S*. *pneumoniae* in the initial 10 base consensus operator and this is notated as “K”; 2) the middle bases of the 10-base sequence, previously notated as “n”, are the same in the distal site (“AA”), but “GT” in the proximal site; and 3) both had three of the four bases of our proposed extended consensus being (<u>R</u>nYKACAnnTGTA<u>R</u>n<u>Y</u>) the opposite purine or pyrimidine base as compared to pneumococcus (Figure 1B). Pneumococcal CopY bound to the distal site implying that the predictions of “K” in the previous 10-base consensus operator and the three changed purines and pyrimidine were correct (Figure 7A). However, pneumococcal CopY did not bind to the proximal site implying that the “AA” notated as “nn” in the previous 10-base consensus sequence indeed needed to be “AA” (Figure 7B).

**Figure 7.**
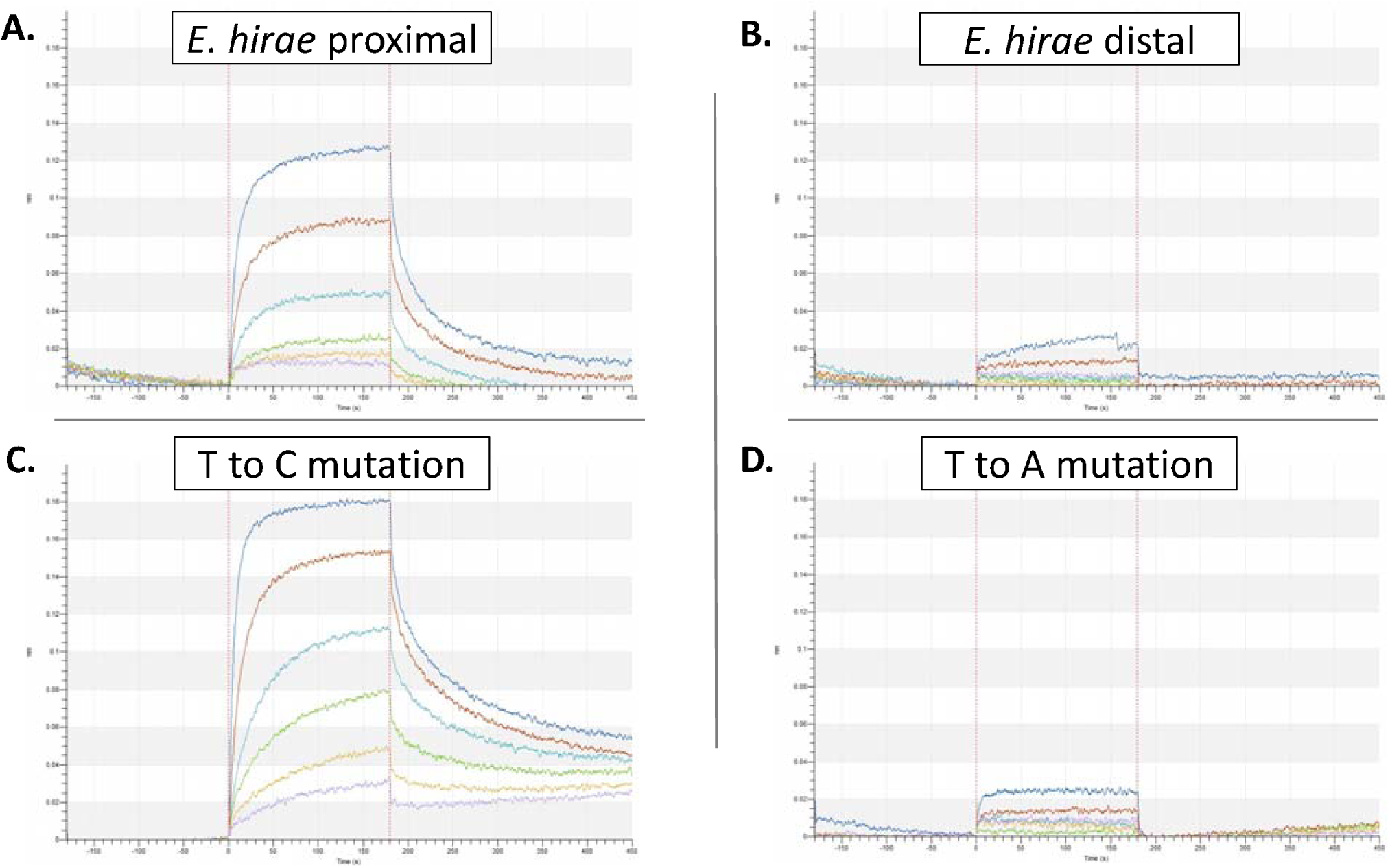
Pneumococcal CopY binds to *E*. *hirae* DNA in accordance with the newly proposed consensus *cop* operon operator. Affinity of CopY binding to various DNA fragments was determined using the following concentrations of CopY 1000 nM (blue), 500 nM (red), 250 nM (light blue), 125 nM (green), 62.5 nM (orange), 15.6 nM (purple) (A-D). **A.** *E*. *hirae* proximal site. **B.** *E*. *hirae* distal site **C.** 19-base fragment with a T to C mutation. **D.** 19-base fragment with a T to A mutation. For each figure, data is representative of three experimental replicates.

Lastly, we tested our hypothesis that it is a pyrimidine or purine in the given positions 1, 3, 14, and 16, and not the specific base that matters by mutating the pyrimidine T at position 3 (ATTGACAAATGTAGAT), to C (pyrimidine) and to A (purine) in the 19-base DNA fragment. We used the 19-base fragment versus the 16-base fragment here to better detect changes in binding affinity between the samples. Of the four bases outside of the 10-base consensus sequence that we made predictions for, the pyrimidine at position 3 is the only base that was the same in the *E*. *hirae* DNA fragment. As expected within our model, the T to C mutation did not significantly alter the binding affinity as compared to the 19-base fragment, while the T to A mutation completely abolished binding (Figure 7C, D). Taken together, we believe the pneumococcal consensus operator is RnYKACAAATGTARnY as opposed to the previously reported KACAnnTGTA (Figure 8).

**Figure 8.**
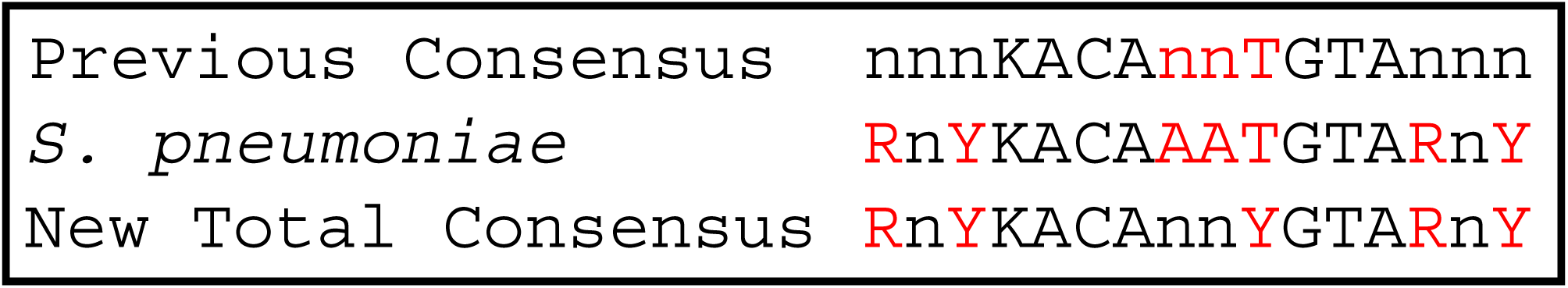
Chart representing the previous, newly proposed pneumococcal, and newly proposed CopY/R protein family consensus *cop* operon operator. Bases that change from the initial 10-base consensus operator are highlighted in red.

## DISCUSSION

Here, we populated a list of pneumococcal CopY homologs, assessed them for the number of upstream operators, determined the affinity of pneumococcal CopY for each of its two operators, and elucidated the complete sequence of the pneumococcal *cop* operon operator (RnYKACAAATGTARnY) which is greatly expanded from the previously reported KACAnnTGTA (Figure 8). As expected from the homology of the *cop* operon repressors, we observed binding of one species’ repressor to the DNA of another, but this binding was not absolute as pneumococcal CopY bound to only one of the *E*. *hirae* sites.

Given the previous 10-base consensus sequence, it was plausible to propose that like in *L*. *lactis*, there were additional binding sites that the *cop* repressor could bind. Furthering the biological relevance of this proposition was that some sequences in *S*. *pneumoniae* corresponded to putative promoter regions of genes and operons upregulated under copper stress, thus implying that CopY was a master regulator of several operons under copper stress (11). By showing no binding of these sequences that followed the 10-base consensus, and the 10-base sequence itself, we demonstrated that the 10-base sequence was not sufficient for binding. This fact ultimately led us to searching for and deriving the pneumococcal consensus sequence. Given this new pneumococcal consensus sequence, a new search in multiple *Streptococcal* species for potential operators yielded no additional sites (49). However, as in the *L*. *lactis* CopR binding to multiple operators in its genome, this is likely not the case with all bacteria containing CopY homologs (48).

In generating the list of CopY/R genes and proteins, we also were able to look upstream of the gene and compile what we believe to be a general consensus operator site which differs slightly from the newly proposed pneumococcal version. The two changes from the pneumococcal operator we propose are 1) returning to the “nn” in the middle of the previous 10-base consensus sequence and 2) changing base 10 from “T” to “Y” based on having T or C in the meme suite alignment. These changes yield what we propose to be the new consensus *cop* operon operator “RnYKACAnnYGTARnY” (Figure 8). This new consensus operator was able to explain why CopY bound a subset of our tested sites. Furthermore, the model also explains the results presented by Magnani *et*. *al*. using the *L*. *lactis* CopR protein (Table 2) (48). As such, consensus operator sequences for various repressors should be revisited to better reveal potential binding interactions within their respective genomes.

Previous studies characterizing pneumococcal CopY were carried out with DNA containing only one intact operator (17, 32, 33). However, we show that *Streptococcus* and many other genera have two binding sites upstream of the *cop* operon. In some cases (e.g., *Streptococcus*), these sites are identical, while other sites have slight sequence variation (e.g., *E*. *hirae)* (Figure 1A). Despite having two identical 21-base repeat operators, the *S*. *pneumoniae cop* operon two-operator DNA does not have tighter binding to CopY as compared to the proximal or distal operator. This result is contrary to what was expected as more DNA-binding sites in sequence tend to increase overall protein affinity for those sites of regulation. While it is clear that CopY does not need a second operator present to bind DNA, this result does not necessarily rule out that *cop* operons with two operators is more tightly regulated than if only one operator was present.

Regarding how stringently the *cop* operon is controlled, we propose that the two pneumococcal operators do indeed serve to add additional restraint to *cop* operon transcription. We hypothesize that the two pneumococcal operators do indeed serve to add additional restraint to *cop* operon transcription. We also suggest that the distal operator prevents sigma factors from binding at the *copY* -35 element. and the proximal operator occludes RNA polymerase; establishing two layers of repression for the *cop* operon (51). This reasoning is also consistent to the proposed hypothesis of why there are two operators in the antibiotic resistance repressor BlaI (52). These hypothesizes are the subject of our group’s ongoing research. Still, *S*. *pneumoniae* with its two CopY operators and with multiple levels of regulation comes as a surprise for an operon in which A) upregulation is linked to increased pneumococcal survival in the host and B) Δ*copY* mutant has increased virulence in mice (27, 53). We anticipate that further study of these systems in *S*. *pneumoniae* will yield clues as to the competitive advantages or selective pressures of the *cop* operon in its pathological context.

## CONFLICTS OF INTEREST

There are no conflicts to declare.

## ACKNOWLEDGEMENTS

The authors would like to thank Rachel Wong, the University of Arizona Functional Genomics Core Facility, and Richard Yip for assistance and support using the Octet Red384. This work was funded by an NIGMS grant 1R35128653 (MDLJ).

## REFERENCES

1. Irving H, Williams RJP. 1953. 637. The stability of transition-metal complexes. Journal of the Chemical Society (Resumed) doi:10.1039/JR9530003192:3192–3210.

2. Imlay JA. The mismetallation of enzymes during oxidative stress.

3. Braymer JJ, Giedroc DP. 2014. Recent developments in copper and zinc homeostasis in bacterial pathogens. Current opinion in chemical biology 19:59–66.

4. Salgado CD, Sepkowitz KA, John JF, Cantey JR, Attaway HH, Freeman KD, Sharpe PAM, Michels HT, Schmidt MG. 2013. Copper Surfaces Reduce the Rate of Healthcare-Acquired Infections in the Intensive Care Unit. Infection Control and Hospital Epidemiology 34:479–486.

5. Warnes SL, Keevil CW. 2016. Lack of Involvement of Fenton Chemistry in Death of Methicillin-Resistant and Methicillin-Sensitive Strains of Staphylococcus aureus and Destruction of Their Genomes on Wet or Dry Copper Alloy Surfaces. Appl Environ Microbiol 82:2132–2136.

6. Macomber L, Hausinger RP. 2011. Mechanisms of nickel toxicity in microorganisms. Metallomics 3:1153–1162.

7. Kumar V, Mishra RK, Kaur G, Dutta D. 2017. Cobalt and nickel impair DNA metabolism by the oxidative stress independent pathway. Metallomics 9:1596–1609.

8. Kehl-Fie TE, Skaar EP. 2010. Nutritional immunity beyond iron: a role for manganese and zinc. Curr Opin Chem Biol 14:218–224.

9. Hood MI, Skaar EP. 2012. Nutritional immunity: transition metals at the pathogen-host interface. Nat Rev Microbiol 10:525–537.

10. Macomber L, Imlay JA. 2009. The iron-sulfur clusters of dehydratases are primary intracellular targets of copper toxicity. Proc Natl Acad Sci U S A 106:8344–8349.

11. Johnson MD, Kehl-Fie TE, Rosch JW. 2015. Copper intoxication inhibits aerobic nucleotide synthesis in Streptococcus pneumoniae. Metallomics 7:786–794.

12. Djoko KY, Phan MD, Peters KM, Walker MJ, Schembri MA, McEwan AG. 2017. Interplay between tolerance mechanisms to copper and acid stress in Escherichia coli. Proc Natl Acad Sci U S A 114:6818–6823.

13. Djoko KY, McEwan AG. 2013. Antimicrobial action of copper is amplified via inhibition of heme biosynthesis. ACS Chem Biol 8:2217–2223.

14. Macomber L, Rensing C, Imlay JA. 2007. Intracellular copper does not catalyze the formation of oxidative DNA damage in Escherichia coli. J Bacteriol 189:1616–1626.

15. Miao X, He J, Zhang L, Zhao X, Ge R, He QY, Sun X. 2018. A Novel Iron Transporter SPD_1590 in Streptococcus pneumoniae Contributing to Bacterial Virulence Properties. Front Microbiol 9:1624.

16. Honsa ES, Johnson MD, Rosch JW. 2013. The roles of transition metals in the physiology and pathogenesis of. Frontiers in cellular and infection microbiology 3:92.

17. Shafeeq S, Yesilkaya H, Kloosterman TG, Narayanan G, Wandel M, Andrew PW, Kuipers OP, Morrissey JA. 2011. The cop operon is required for copper homeostasis and contributes to virulence in Streptococcus pneumoniae. Molecular microbiology 81:1255–1270.

18. Rosch JW, Sublett J, Gao G, Wang YD, Tuomanen EI. 2008. Calcium efflux is essential for bacterial survival in the eukaryotic host. Molecular microbiology 70:435–444.

19. Rosch JW, Gao G, Ridout G, Wang YD, Tuomanen EI. 2009. Role of the manganese efflux system mntE for signalling and pathogenesis in Streptococcus pneumoniae. Molecular microbiology 72:12–25.

20. Honsa ES, Johnson MDL, Rosch JW. 2013. The roles of transition metals in the physiology and pathogenesis of Streptococcus pneumoniae. Frontiers in Cellular and Infection Microbiology 3.

21. Shafeeq S, Kuipers OP, Kloosterman TG. 2013. The role of zinc in the interplay between pathogenic streptococci and their hosts. Molecular microbiology 88:1047–1057.

22. Strausak D, Solioz M. 1997. CopY is a copper-inducible repressor of the Enterococcus hirae copper ATPases. The Journal of biological chemistry 272:8932–8936.

23. Smaldone GT, Helmann JD. 2007. CsoR regulates the copper efflux operon copZA in Bacillus subtilis. Microbiology 153:4123–4128.

24. Corbett D, Schuler S, Glenn S, Andrew PW, Cavet JS, Roberts IS. 2011. The combined actions of the copper-responsive repressor CsoR and copper-metallochaperone CopZ modulate CopA-mediated copper efflux in the intracellular pathogen Listeria monocytogenes. Molecular microbiology 81:457–472.

25. Vollmecke C, Drees SL, Reimann J, Albers SV, Lubben M. 2012. The ATPases CopA and CopB both contribute to copper resistance of the thermoacidophilic archaeon Sulfolobus solfataricus. Microbiology 158:1622–1633.

26. Jacobs AD, Chang FMJ, Morrison L, Dilger JM, Wysocki VH, Clemmer DE, Giedroc DP. 2015. Resolution of Stepwise Cooperativities of Copper Binding by the Homotetrameric Copper-Sensitive Operon Repressor (CsoR): Impact on Structure and Stability. Angewandte Chemie-International Edition 54:12795–12799.

27. Johnson MD, Kehl-Fie TE, Klein R, Kelly J, Burnham C, Mann B, Rosch JW. 2015. Role of copper efflux in pneumococcal pathogenesis and resistance to macrophage-mediated immune clearance. Infect Immun 83:1684–1694.

28. Singh K, Senadheera DB, Levesque CM, Cvitkovitch DG. 2015. The copYAZ Operon Functions in Copper Efflux, Biofilm Formation, Genetic Transformation, and Stress Tolerance in Streptococcus mutans. J Bacteriol 197:2545–2557.

29. Arguello JM, Gonzalez-Guerrero M, Raimunda D. 2011. Bacterial transition metal P(1B)-ATPases: transport mechanism and roles in virulence. Biochemistry 50:9940–9949.

30. Hava DL, Camilli A. 2002. Large-scale identification of serotype 4 Streptococcus pneumoniae virulence factors. Molecular microbiology 45:1389–1406.

31. Stoyanov JV, Brown NL. 2003. The Escherichia coli copper-responsive copA promoter is activated by gold. The Journal of biological chemistry 278:1407–1410.

32. Neubert MJ, Dahlmann EA, Ambrose A, Johnson MDL. 2017. Copper Chaperone CupA and Zinc Control CopY Regulation of the Pneumococcal cop Operon. Msphere 2.

33. Glauninger H, Zhang Y, Higgins KA, Jacobs AD, Martin JE, Fu Y, Coyne Rd HJ, Bruce KE, Maroney MJ, Clemmer DE, Capdevila DA, Giedroc DP. 2018. Metal-dependent allosteric activation and inhibition on the same molecular scaffold: the copper sensor CopY from Streptococcus pneumoniae. Chem Sci 9:105–118.

34. Liu T, Ramesh A, Ma Z, Ward SK, Zhang LM, George GN, Talaat AM, Sacchettini JC, Giedroc DP. 2007. CsoR is a novel Mycobacterium tuberculosis copper-sensing transcriptional regulator. Nature Chemical Biology 3:60–68.

35. Fu Y, Tsui HC, Bruce KE, Sham LT, Higgins KA, Lisher JP, Kazmierczak KM, Maroney MJ, Dann CE, 3rd, Winkler ME, Giedroc DP. 2013. A new structural paradigm in copper resistance in Streptococcus pneumoniae. Nat Chem Biol 9:177–183.

36. Fu Y, Bruce KE, Wu HW, Giedroc DP. 2016. The S2 Cu(I) site in CupA from Streptococcus pneumoniae is required for cellular copper resistance. Metallomics 8:61–70.

37. Changela A, Chen K, Xue Y, Holschen J, Outten CE, O’Halloran TV, Mondragon A. 2003. Molecular basis of metal-ion selectivity and zeptomolar sensitivity by CueR. Science 301:1383–1387.

38. Safo MK, Zhao QX, Ko TP, Musayev FN, Robinson H, Scarsdale N, Wang AHJ, Archer GL. 2005. Crystal structures of the BlaI repressor from Staphylococcus aureus and its complex with DNA: Insights into transcriptional regulation of the bla and mec operons. Journal of Bacteriology 187:1833–1844.

39. Arede P, Oliveira DC. 2013. Proteolysis of mecA repressor is essential for expression of methicillin resistance by Staphylococcus aureus. Antimicrob Agents Chemother 57:2001–2002.

40. Altschul SF, Gish W, Miller W, Myers EW, Lipman DJ. 1990. Basic local alignment search tool. J Mol Biol 215:403–410.

41. Stols L, Gu MY, Dieckman L, Raffen R, Collart FR, Donnelly MI. 2002. A new vector for high-throughput, ligation-independent cloning encoding a tobacco etch virus protease cleavage site. Protein Expression and Purification 25:8–15.

42. Portmann R, Magnani D, Stoyanov JV, Schmechel A, Multhaup G, Solioz M. 2004. Interaction kinetics of the copper-responsive CopY repressor with the cop promoter of Enterococcus hirae. J Biol Inorg Chem 9:396–402.

43. Collins TC, Dameron CT. 2012. Dissecting the dimerization motif of Enterococcus hirae’s Zn(II)CopY. Journal of Biological Inorganic Chemistry 17:1063–1070.

44. Gupta RS, Lo B, Son J. 2018. Phylogenomics and Comparative Genomic Studies Robustly Support Division of the Genus Mycobacterium into an Emended Genus Mycobacterium and Four Novel Genera. Front Microbiol 9:67.

45. Bailey TL, Johnson J, Grant CE, Noble WS. 2015. The MEME Suite. Nucleic Acids Res 43:W39–49.

46. Bailey TL, Boden M, Buske FA, Frith M, Grant CE, Clementi L, Ren J, Li WW, Noble WS. 2009. MEME SUITE: tools for motif discovery and searching. Nucleic Acids Res 37:W202–208.

47. Bailey TL, Elkan C. 1994. Fitting a mixture model by expectation maximization to discover motifs in biopolymers. Proc Int Conf Intell Syst Mol Biol 2:28–36.

48. Magnani D, Barre O, Gerber SD, Solioz M. 2008. Characterization of the CopR regulon of Lactococcus lactis IL1403. J Bacteriol 190:536–545.

49. Mrazek J, Xie S. 2006. Pattern locator: a new tool for finding local sequence patterns in genomic DNA sequences. Bioinformatics 22:3099–3100.

50. Odermatt A, Solioz M. 1995. Two trans-acting metalloregulatory proteins controlling expression of the copper-ATPases of Enterococcus hirae. J Biol Chem 270:4349–4354.

51. Slager J, Aprianto R, Veening JW. 2018. Deep genome annotation of the opportunistic human pathogen Streptococcus pneumoniae D39. Nucleic Acids Res 46:9971–9989.

52. Gregory PD, Lewis RA, Curnock SP, Dyke KG. 1997. Studies of the repressor (BlaI) of beta-lactamase synthesis in Staphylococcus aureus. Mol Microbiol 24:1025–1037.

53. van Opijnen T, Camilli A. 2012. A fine scale phenotype-genotype virulence map of a bacterial pathogen. Genome Res 22:2541–2551.

