## Supplemental Table 1 and Figures 1-3 for "Characterization of consensus operator site for *Streptococcus pneumoniae* copper repressor, CopY"

Supplemental information

**Supplementary Figure Legends**

**Supplementary Table 1. Primer Table.**

**Supplementary Table 2. Homologous protein sequences to *S. pneumoniae* TIRG4 CopY from BLAST search.**

**Supplementary Table 3. Homologous protein and DNA sequences to *S. pneumoniae* TIRG4 CopY with unique protein accession numbers.**

**Supplementary Table 4. Homologous protein and DNA sequences to *S. pneumoniae* TIRG4 CopY with unique genome accession numbers.**

**Supplementary Table 5. CopY consensus *cop* operon operators from MEME suite from unique genome accession numbers.**

**Supplementary Table 6. CopY consensus *cop* operon operators from Table S4 with only one *cop* operon operator.**

**Supplementary Table 7. Organisms from Table S4 with no consensus *cop* operon operator.**

**Supplemental Table 1. List of primers used for BLI.** Each of the primers in the table have a 5’ biotinylation and are the forward primer used in the dsDNA for BLI assay. Complementary reverse primers for each fragment were used to generate the dsDNA (not shown).

**Supplemental Figure 1. Distances in bases between operators in species that had two operators.** A python script was generated to count and plot on a graph the number of bases between operators (available at <https://github.com/Van-Doorslaer/Alvin_et_al_2018>).

**Supplemental Figure 2. CopY does not bind predicted sites continued.** CopY at 3 µM was used to assess binding to DNA fragments containing potential CopY operators upstream of the respective genes *SP_1863* (red) and *SP_2073* (light blue), with controls, two-site DNA (blue, positive control) and scram (green, negative control).

**Supplemental 3. Affinity of CopY for 16bp fragment**. Binding affinity of CopY to the 16-base fragment was determined using the following concentrations of protein 1000 nM (blue), 500 nM (red), 250 nM (light blue), 125 nM (green), 62.5 nM (orange), 15.6nM (purple).

**Supplemental Table 1**

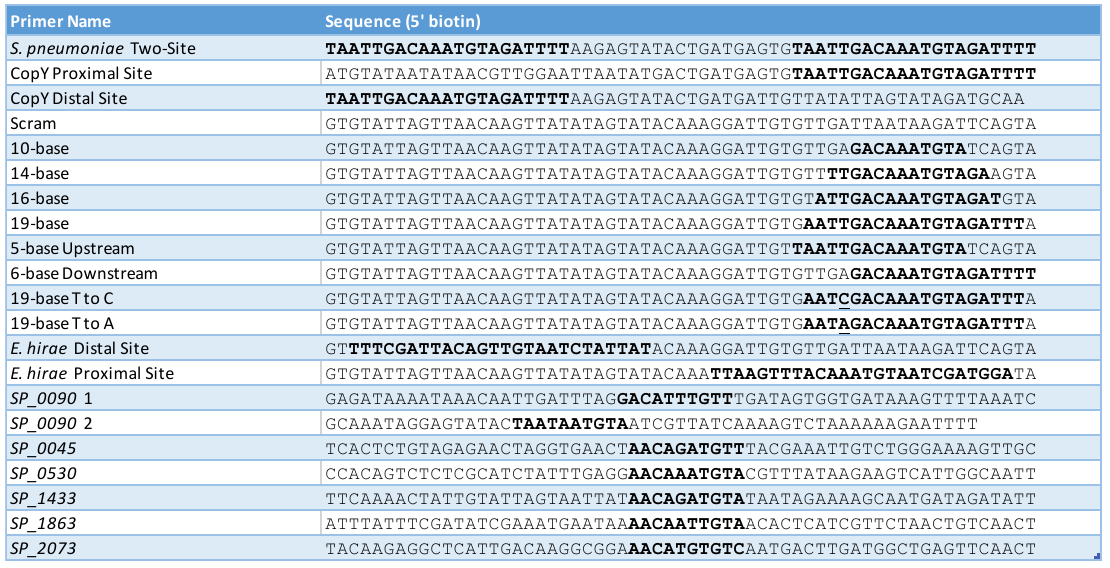

**Supplemental Figure 1**

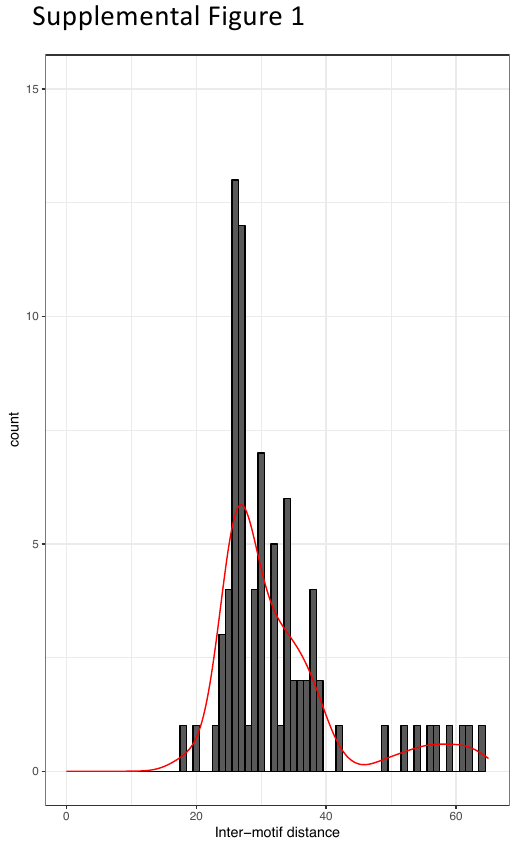

**Supplemental Figure 2**

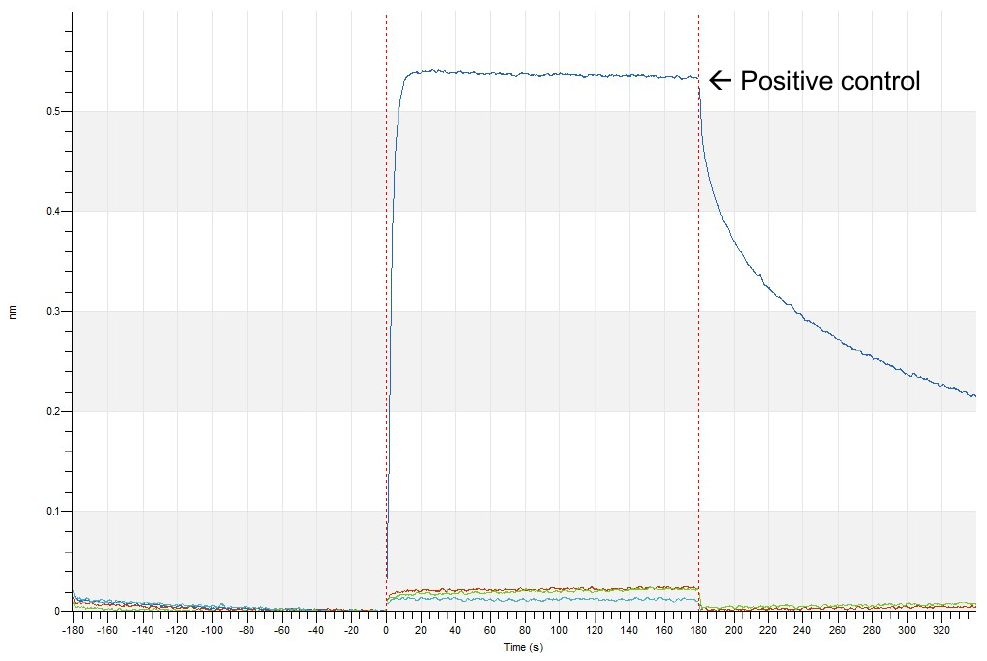

Supplemental Figure 3

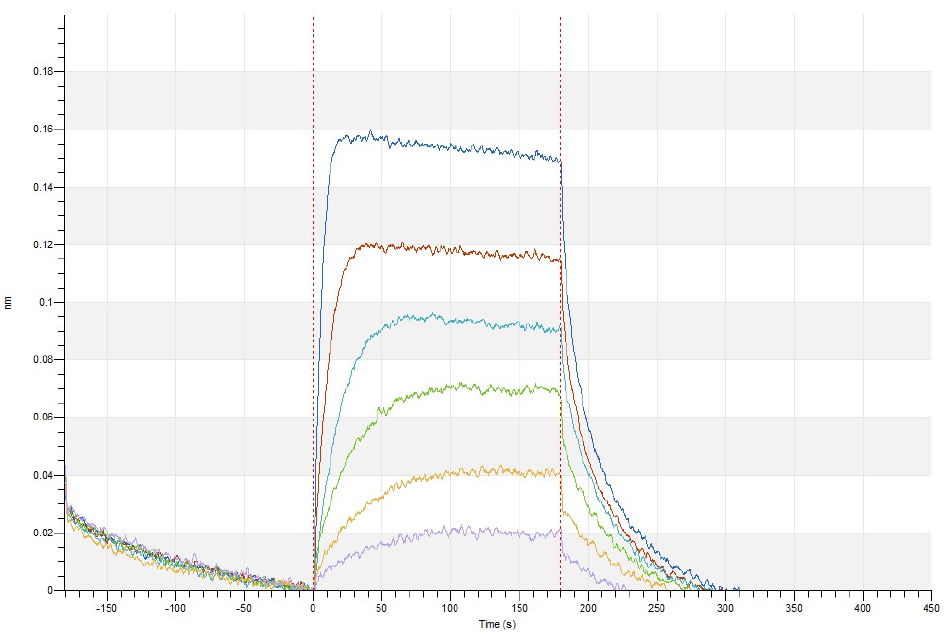

Supplemental Figure 4

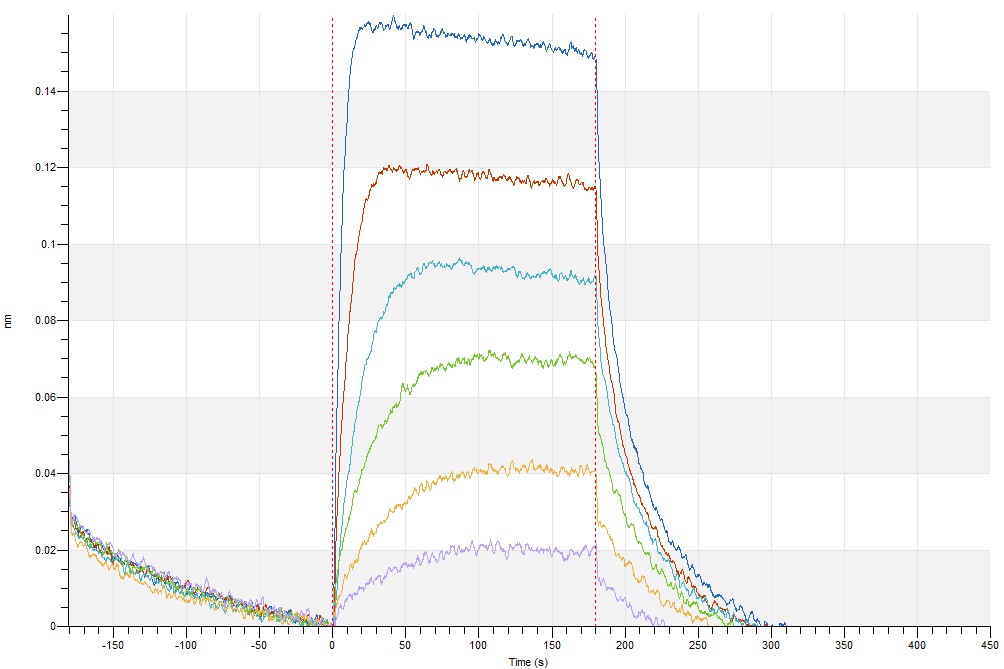
